# A probabilistic view of forbidden links: their prevalence and their consequences for the robustness of plant-hummingbird communities

**DOI:** 10.1101/2024.07.04.602032

**Authors:** François Duchenne, Elisa Barreto, Esteban A. Guevara, Holger Beck, Carolina Bello, Rafaela Bobato, Daniela Bôlla, Emanuel Brenes, Nicole Büttner, Ana P. Caron, Nelson Chaves-Elizondo, María J. Gavilanes, Alejandro Restrepo-González, Jose Alejandro Castro, Miriam Kaehler, Tiago Machado-de-Souza, Miguel Machnicki-Reis, Andrés Sebastián F. Marcayata, Cauã G. de Menezes, Andrea Nieto, Rafael de Oliveira, Ricardo A. C. de Oliveira, Friederike Richter, Bryan G. Rojas, Luciele L. Romanowski, Wellinton L.de Souza, Danila S. Veluza, Ben Weinstein, Rafael Wüest, Thais B. Zanata, Krystal Zuniga, María A. Maglianesi, Tatiana Santander, Isabella G. Varassin, Catherine H. Graham

**Author notes:** **<u>Data availability statement:</u>** The data will be available on Zenodo upon manuscript acceptance. R codes used for analyses are available here: https://github.com/f-duchenne/EPHI_paper. **<u>Competing Interest Statement:</u>** The authors declare no competing interests. **<u>Author contributions:</u>** FD designed the study, with inputs from ElB and CHG, and performed all analyses. CHG was responsible for overall project design and coordination across the three countries. IGV, TS and MM were responsible for country level scientific and logistic coordination/data collection with major assistance from TM-S, TBZ, AN, FR, and EBr. HB, RB, DB, EBr, NB, APC, NCE, ARG, JAC, MK, TM-S, MM-R, CGM, ASMF, AN, RO, FR, BGR, LLR, WLS, DSV, and KZ collected the data. EBa, FD, CB, EBr, MK, TS, TM-S, RW, BW, and IGV curated the data. FD wrote the first draft and all authors contributed to the final writing and editing.

## Abstract

The presence in ecological communities of unfeasible species interactions, termed forbidden links, due to physiological or morphological exploitation barriers has been long debated, but little direct evidence has been found. Forbidden links are likely to make ecological communities less robust to species extinctions, stressing the need to assess their prevalence. Here, we used a dataset of plant-hummingbird interactions, coupled with a Bayesian hierarchical model, to assess the importance of exploitation barriers in determining species interactions. We found evidence for exploitation barriers between flowers and hummingbirds across the 32 studied communities, however, the proportion of forbidden links changed drastically among communities, because of changes in trait distributions. The higher the proportion of forbidden links, the more they decreased network robustness, because of constraints on interaction rewiring. Our results suggest that exploitation barriers are not rare in plant-hummingbird communities and have the potential to limit the rescue of species experiencing partner extinction.

## Introduction

Identifying the factors influencing species interactions is a major goal of ecology, because the organization of species interactions shapes the stability of natural communities (Bascompte 2009; Kéfi *et al*. 2016; Thébault & Fontaine 2010). Ecologists have investigated the mechanisms that determine species interactions, henceforth linkage rules, highlighting that interactions are often structured by species abundances and morphological traits rather than species identity (Bartomeus *et al*. 2016; Gravel *et al*. 2013; Machado-de-Souza *et al*. 2019; Vázquez *et al*. 2009b, a). Given that rules are associated with traits and abundance, species can interact with different sets of species through space and time (CaraDonna *et al*. 2017; Tylianakis & Morris 2017), which makes communities less vulnerable to the loss of specific species (Kaiser-Bunbury *et al*. 2010; Ramos-Jiliberto *et al*. 2012; Vizentin-Bugoni *et al*. 2020). To date, much progress has been made on identifying rules that influence the presence of species interactions but not what causes the absence of interactions. While the latter is simply the other side of the same coin, a focus on what influences the absence of interactions might highlight constraints on the distribution and flexibility of interspecific interactions that are otherwise overlooked.

Ecological networks are often very sparse, meaning that the number of observed interactions is often much lower than the number of possible interactions (Busiello *et al*. 2017; Jordano 2016). A few explanations have been proposed to explain the sparsity of ecological networks: insufficient sampling where some interactions are missed (Doré *et al*. 2021; Jordano 2016), low species abundances (Vázquez *et al*. 2009a, b) or trait matching constraints (Santamaría & Rodríguez-Gironés 2007). Trait matching often considers trait complementarity and barrier (Santamaría & Rodríguez-Gironés 2007). Trait complementarity is a continuous measure, such as similarity between corolla length and proboscis or bill length (Gonzalez & Loiselle 2016; Junker *et al*. 2013; Sonne *et al*. 2020; Vázquez *et al*. 2009a). In contrast, trait barrier refers to compatible *versus* incompatible traits, which lead to an on/off mechanism, called an exploitation barrier (Stang *et al*. 2009). Among the many missing interactions in a network, only the ones due to exploitation barriers are forbidden links (*i*.*e*. links invariantly absent). The proportion of these forbidden links is important for understanding spatio-temporal dynamics of communities, because they are links that would be absent regardless of the context, such as species abundances, or sampling frequency.

In case of species extinctions, forbidden links might limit species’ ability to change interaction partners, referred to as rewiring, and thus could be critical for understanding and assessing community robustness. Robustness is often assessed by simulating species extinctions and evaluating how fast a given network collapses (Memmott *et al*. 2004; Pocock *et al*. 2012). A robust network is a network that needs a high number of primary extinctions to collapse. Interaction rewiring is a key mechanism influencing network robustness (Kaiser-Bunbury *et al*. 2010; Ramos-Jiliberto *et al*. 2012; Vizentin-Bugoni *et al*. 2020). The more species can rewire when their interactions are lost due to primary extinctions, the higher the robustness. Knowledge of the prevalence of forbidden links in the network would provide insights into the robustness of the interaction networks to perturbation.

Evidencing a forbidden link requires demonstrating that the absence of interaction is not due to insufficient sampling or stochasticity, but to an exploitation barrier. So far, forbidden links have been identified mainly using indirect evidence by looking at network properties, such as the distribution of interaction frequencies (Eklöf *et al*. 2013; Jordano *et al*. 2003) and topological characteristics of interaction networks (Santamaría & Rodríguez-Gironés 2007). This indirect evidence does not explicitly link the forbidden links with their causal mechanisms, but see Olesen *et al*. (2010). To obtain direct evidence of forbidden links, we need to investigate the mechanisms driving the absence of interactions between pairs of species. However, statistical models simultaneously testing different trait matching linkage rules in empirical mutualistic networks are scarce (but see Stang *et al*. 2009). Studies often either consider trait complementarity (Klumpers *et al*. 2019; Sonne *et al*. 2020; Zhao *et al*. 2022) or exploitation barrier (Vizentin-Bugoni *et al*. 2014) or both rules but in separate models (Sazatornil *et al*. 2016). In addition, the concept of forbidden links has been criticized because studies often neglect the intraspecific trait variability that might reduce their prevalence (González-Varo & Traveset 2016). Overcoming these limitations requires repeated sampling across multiple species and individuals from the same species and a probabilistic view of species interactions (Song *et al*. 2020) to measure forbidden links.

To assess how linkage rules and trait distributions determine the sparsity of ecological networks and their robustness, we used plant-hummingbird interaction data collected over three different elevation gradients from different biogeographic regions (Dalsgaard *et al*. 2021; McGuire *et al*. 2014). These regions include the mountains in the Atlantic Forest, in Brazil, the Tropical Andes, in Ecuador, and the Talamanca Mountains, in Costa Rica. We developed a mechanistic and hierarchical Bayesian model considering two linkage rules: possible or forbidden links due to exploitation barrier and a gradient of interaction frequencies due to trait complementarity. Then, merging estimated linkage rules and the observed variation in trait distribution over elevation, we used simulations to estimate how network robustness to plant extinction depended on the linkage rules (Fig. 1). We show that linkage rules and trait distributions together drive the robustness of plant-hummingbird communities to species extinctions.

**Figure 1:**
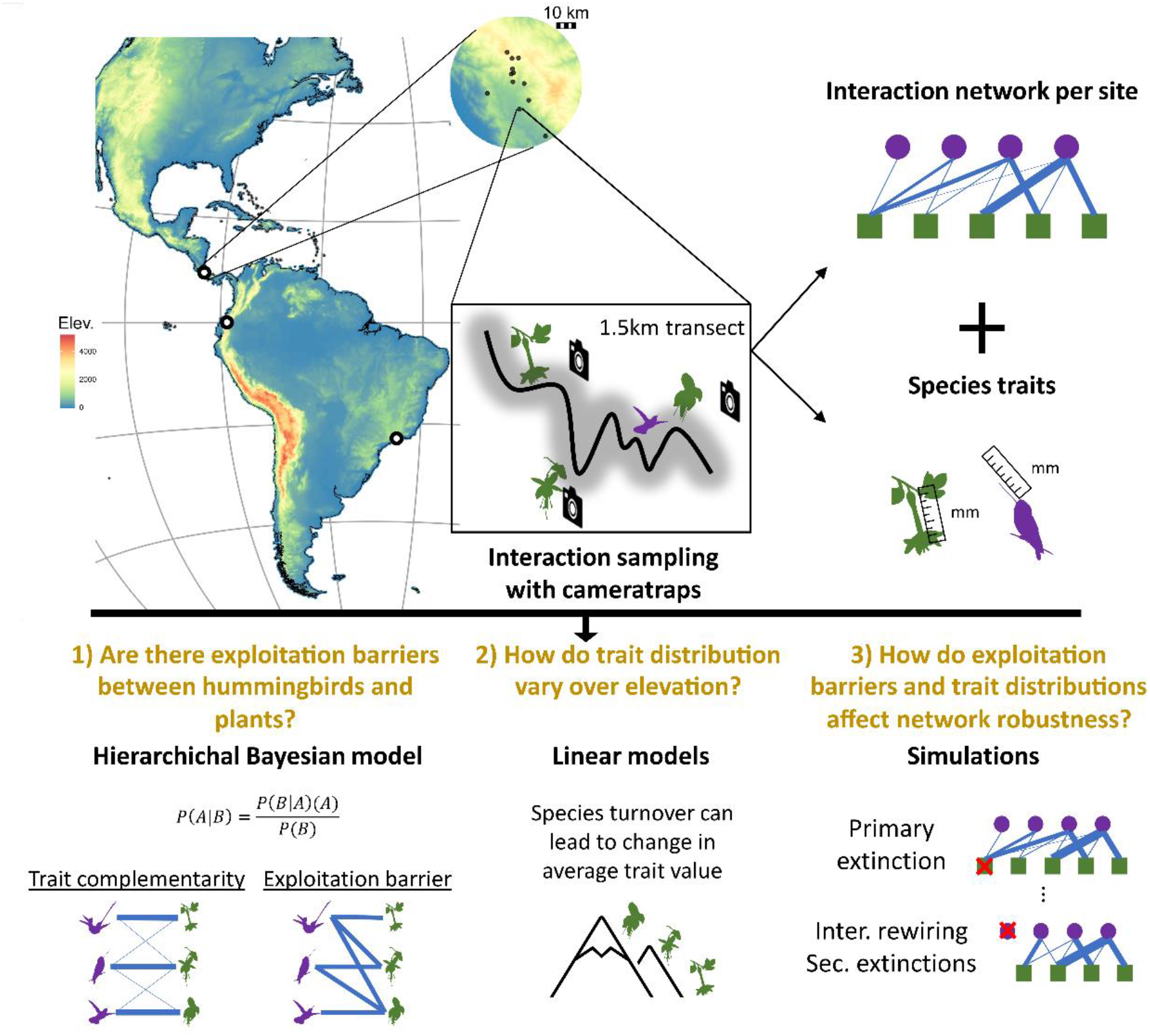
Schematic representation of the methodological steps to investigate drivers of sparsity and their effects on network robustness. On the map, the white points show the location of the three regions, while the zoomed map shows an example of one region, 12 study sites in Costa Rica and a hypothetical transect in one study site. The right part of the figure describes data collection that was done in the transect in a schematic way. Research questions are highlighted below in yellow, while bold black titles represent the methodological steps followed to answer the questions. Elev. = elevation (in meters above sea level); Inter. rewiring = interaction rewiring; Sec. extinctions = secondary extinctions.

## Methods

### Interaction dataset

We collected data in 32 sites located in Brazil, Costa Rica and Ecuador, between 10 and 3500 meters above sea level. Each site was sampled during about two years, spread between February 2017 and December 2022 (*cf*. Table S1, Fig. 1). Each site consisted of one 1.5 km transect along which we tracked plant-hummingbird interactions by installing 12 time-lapse cameras at ∼1 meter distance from open flowers, up to 2 m above the ground, during ∼3 days each month. The cameras took an image every second from dawn to dusk (∼ 12 h), generating ∼ 43,200 images per camera per day. Images were analyzed with DeepMeerkat software (Weinstein 2018) which returned candidate frames with hummingbirds. These candidate frames were reviewed manually to identify birds to the species level and determine pairwise plant-hummingbird interactions. In parallel, every month, a flower census was done on each transect to estimate the abundance of available flowers (*cf*. Supplementary Methods).

The overall dataset includes 80,988 interactions detected between 79 hummingbird species and 640 plant species. Since we are interested in linkage rules of mutualistic interactions, we excluded 5,678 interactions in which hummingbirds were observed robbing nectar instead of entering the flower by the corolla opening. These interactions are likely not mutualistic because when piercing, hummingbirds do not contact the reproductive organs of the flower, and thus do not contribute to pollination. Furthermore, since we are interested in trait complementarity and exploitation barrier, we used only the subset of 441 plant species and 73 hummingbird species for which we have trait measures, corresponding to 69,201 interactions. Countries exhibited strong differences in species richness. The studied sites in Ecuador were three times more diverse than those in Brazil (Fig. S1), thus we decided to fit one model per country.

For each camera and each hummingbird species, we summed interactions recorded with the focal plant. When no interaction was detected with a hummingbird species that was known to occur at that site (at least one recorded interaction along the transect over the entire sampling period), the observed interaction count was set to zero. We then modelled this observed interaction count to disentangle the mechanisms producing these zeros (*cf*. below).

### Morphological traits of plants and hummingbirds

To calculate trait matching between a hummingbird and a plant, we used the corolla length for plant flowers and the bill length of hummingbirds measured in the field and compiled from the literature (cf. Supplementary Methods).

The signal of trait matching between corolla and bill length in determining interaction frequencies could likely be blurred by: (i) the fact that hummingbird protrude their tongues beyond the tips of their bill to reach nectar allowing them to interact with flowers with a corolla longer than their bill (Paton & Collins 1989), and (ii) trait matching is not necessarily the rule in mutualism, coevolution between plant and pollinators can turn in an arms race maintaining trait mismatch between interacting species (Paudel *et al*. 2016; Week & Nuismer 2021). To deal with these potential issues, we used the bill length to inform two latent variables for each hummingbird species, one modelling the preferred corolla length and the other modelling the barrier threshold, the maximum corolla length a flower can have for a given bird to interact with (cf. below).

### Inferring linkage rules

To assess linkage rules, we developed a hierarchical Bayesian model including all species and sites together, correcting for spatial and temporal autocorrelation. In an interaction dataset, non-observed interactions (zeros) can be due to three statistical reasons: (i) non-detection due to insufficient sampling pressure; (ii) overdispersion in the distribution of interaction frequencies that can lead to over-representation of zero; (iii) additional zero-inflation in the distribution of interactions.

To distinguish these three statistical sources of zeros, we modelled interaction frequency as a mixture of Bernoulli process (forbidden/possible links) and a negative-binomial distribution accounting for the sampling duration. Our model thus encompassed two nested submodels, equations (1) and (2), modelling zero-inflation and interaction counts, respectively. The submodel of zero-inflation (Bernouilli process), called the exploitation barrier submodel, was divided in two parts: an intercept (*β*_0_) capturing zero-inflation due to unknown processes and an effect of the exploitation barrier (*β*_1_) capturing the zero-inflation due to forbidden links. The submodel of interaction counts, called the trait complementarity submodel, modelled the number of observed interactions as a function of trait complementarity (*β*_3_), hummingbird phenology (*β*_4_) and sampling duration (log (*D*_*c*_)). We also included random effects accounting for plant effect (*θ*_*j*_), site effect (*θ*_*s*_), temporal variations (*θ*_*sv*_) and hummingbird abundances (*θ*_*si*_). For interaction counts, we used a negative binomial distribution because it deals well with overdispersed data, typical of sparse interaction networks.

For each country, the model is described by the following equations.

**Exploitation barrier submodel**

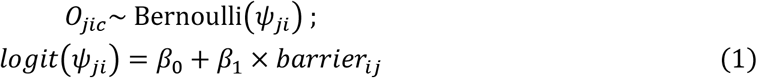

**Trait complementarity submodel**

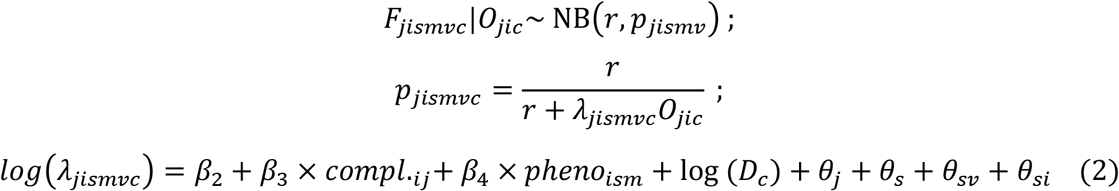

**Latent variables**

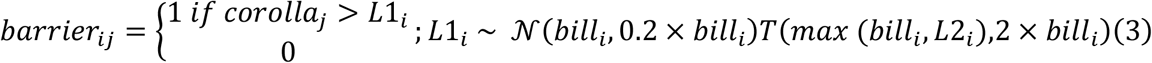

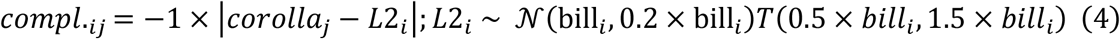

First, in equation (1), we modelled the binary ability of plant species *j* and hummingbird *i* to interact in front of camera *c* (*O*_*jic*_), as a function of exploitation barrier. We used a generalized linear model (GLM) with a binomial error structure and a logit link function (equation (1)), which models the effect of exploitation barrier (*β*_1_) on the probability of interaction (*ψ*_*ji*_) between plant species *j* and hummingbird *i*. Since our measure of exploitation barrier is done at the camera level, it includes a stochastic variability that can be caused by intra-specific trait variability in plants (González-Varo & Traveset 2016). Thus, our estimated probability of interaction indirectly included intraspecific variability in trait values.

Second, in equation (2), we modelled the number of interactions between each plant species *j* and hummingbird *i* on site *s*, month *m*, date *v* and camera *c* (*F*_*jistmc*_), conditionally to the ability of both species to interact. We used a GLM with a negative binomial error structure (with *r* the target for number of successful trials and *p*_*jistm*_ the probability of success in each trial) and a log link function. The average interaction frequencies (*λ*_*jismv*_) are modelled as a function of a general intercept (*β*_2_), the effect of trait complementarity between the plant and the hummingbird (*β*_3_ × *compl*._*ij*_), the hummingbird phenology (*β*_4_ × *pheno*_*im*_, cf. Supplementary methods). Finally, the sampling duration, in hours of daylight (log (*D*_*c*_)), was included as an offset which means that its effect was not estimated but fixed to one. This offset modelled accumulation curve that does not saturate with time, because hummingbirds constantly need to visit flowers. This results in the fact that even if our response variable was interaction count, we corrected by sampling time and thus modelled a number of interactions per hour of sampling.

This submodel also included random effects: a plant species random effect 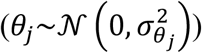 to account for heterogeneity in plant species attractiveness for hummingbirds and a site random effect to account for heterogeneity among sites. To account for spatial autocorrelation among sites we structured the variance-covariance of this random effect using the Euclidean distance among sites: 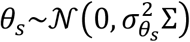 where Σ is the spatial variance-covariance matrix among sites. This matrix was filled with 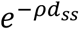 where *ρ* was the scale parameter governing the distance-dependent decay of the spatial correlation and *d*_*ss*_ was the pairwise distance between sites, scaled to be one at the maximum distance. We also included a date (month-year) random effect, modelled as a temporal random walk of order one for each site, to account for temporal autocorrelation: 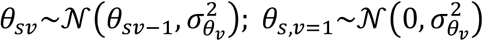. Finally, we used a hummingbird random effect for each site, to account for the fact that the abundance of hummingbirds varies over space 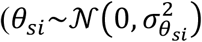.

Latent thresholds (*L*1_*i*_) were estimated for the exploitation barrier for each hummingbird species. This threshold represents the corolla length above which the hummingbird species is not able to interact. To inform the model and accelerate convergence, we used an informative prior, drawing this latent threshold from a Gaussian distribution centred on bill length with standard deviation of 20% of the bill length. We truncated this prior distribution from left and right sides: from half the bill size because we assume that optimal corolla length should not be too far from bill length, and at twice the bill length value, because empirical measures show that tongues are unlikely to extend by more than twice the bill length (Paton & Collins 1989).

The second latent variable (*L*2_*i*_), also estimated for each hummingbird species, corresponds to the optimal corolla length (*i*.*e*. maximizing its interaction frequencies) for the focal bird. Similarly, we used an informative prior, drawing this latent variable from a Gaussian distribution centred on the bill length with standard deviation of 20% the bill length, but truncated to keep values only between the half of bill length and a one and half times the bill length, to avoid unrealistic values. The estimated values of *L*1_*i*_ and *L*2_*i*_ are displayed in Figure S2.

All parameters were estimated simultaneously using Markov Chain Monte Carlo simulations, as implemented in *R2jags* R package (Su & Yajima 2012), with 3 chains, 750,000 iterations with a burnin of 650,000 and a thin rate of 5, which was enough to reach convergence for each of the three countries (≥99% of parameters with Rhat<1.1). Priors are detailed in Supplementary methods.

### Percentage of zeros generated by each linkage rule

To understand how the linkage rules contributed to the sparsity of the networks, we evaluated the percentage of non-interactions (relative to the number of possible interactions) by predicting the percentage of zeros generated by each linkage rule (trait complementarity, exploitation barrier and unknown linkage rule) using the Bayesian model described above (cf.

Supplementary Methods). Since the percentage of zeros generated by trait complementarity depended on the sampling duration, we estimated the percentage of zeros generated by each linkage rule as a function of sampling duration.

### Trait distribution over elevation gradient

To study how species traits vary across elevation, we used a linear mixed-effect model regressing the mean trait value (corolla or bill length) for species detected in a site s as a function of the elevation of site s. We modelled one elevation trend per guild per country by including a triple interaction between elevation, the guild of the species (plant or bird) and the country. We also included the site as a random effect on the intercept.

To study the effect of variation in trait values across elevation on the structure of the networks, we assessed the proportion of forbidden links as the average of 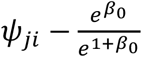 across all pairs of plants and hummingbirds present at a given site. We studied the proportion of forbidden links across elevation using a GLM with a quasi-binomial error structure instead, including elevation in interaction with the country as the response variables.

### Network robustness

To assess network robustness, we used a measure based on coextinction rates, which is the area below the attack tolerance curve (Albert *et al*. 2000). This curve represents the fraction of persisting species of one trophic level, hummingbirds here, as a function of the eliminated species from the other trophic level, plants (Memmott *et al*. 2004; Vizentin-Bugoni *et al*. 2020).

For each site, we used our statistical model to predict initial interaction networks normalized by sampling duration. In these initial networks, pairwise interactions were measured as an average number of visits per day (across the two years of sampling, *cf*. Supplementary Methods). Then, we simulated sequential plant extinctions, and after each primary extinction we calculated the probability of secondary extinction for each hummingbird species (*cf*. Supplementary Methods). The probability of extinction of each hummingbird species depended on the proportion of interactions lost after rewiring. A hummingbird could rewire the interactions lost because of primary plant extinctions if the remaining plant species had compatible traits. To test the effects of forbidden links on robustness we performed simulations with two different cases of rewiring: constrained by trait complementarity only or constrained by trait complementarity and exploitation barriers (*cf*. Supplementary Methods). To explore the dependence of our results on the sequence of extinctions and strength of rewiring, we also performed simulations for two different scenarios of plant extinctions, removing plant species from the more generalists to the more specialists or the opposite, and for three different values of rewiring constraint, *α* ∈ [0,1,2], where α = 0 corresponds to a complete rewiring if no zero-inflation, and α = 2 corresponds to a rewiring strongly constrained by trait matching (*cf*. Supplementary Methods).

We performed 500 simulations for each case (with or without forbidden links, plant extinction scenario, value of rewiring rate and site), and for each we measured the area under the attack tolerance curve. This area is bounded between 0, lowest robustness possible, and 1, highest robustness possible (asymptotic limit).

### Statistical link between forbidden links and robustness

The effect of forbidden links on network robustness was assessed with a GLM with a beta distribution error structure and a *logit* link function, with robustness as the response variable and three discrete explanatory variables in interaction: the discrete variable encoding the presence or absence of forbidden links, the site and the strength of rewiring constraint (*α*). This approach allowed us to test for the effect of forbidden links on robustness for each site, and to test if this effect varied according to easiness of the rewiring. In the case of rewiring-mediated effect of forbidden links on robustness, we expected that the effect of forbidden links on robustness would be stronger when rewiring was high (low constraint) but weak when rewiring was low (high constraint). We ran one GLM for each scenario (plant extinction scenario, specialists or generalists first).

We also tested for significant changes in initial structure when including forbidden links. For that, we used two metrics, connectance and nestedness (cf. Supplementary methods for calculation), and Wilcoxon tests for paired samples.

## Results

We found evidence for both kinds of trait matching, namely trait complementarity and exploitation barriers, with similar patterns across countries (Fig. 2a-b). Interaction frequencies depended on the morphological match between flower corolla length and hummingbird bill length (Fig. 2a & Fig. S2), suggesting that trait complementarity influences plant-hummingbird interactions. On top of trait complementarity, we found a significant exploitation barrier effect in plant-hummingbird interactions (Fig. 2b). This exploitation barrier is evidenced by the strong decrease in the probability of interaction between plants and hummingbirds if the corolla is too long relative to the hummingbird’s bill. This result provides evidence for forbidden links: when there was an exploitation barrier, the probability of observing an interaction dropped, while the probability of observing an invariant zero increased.

**Figure 2:**
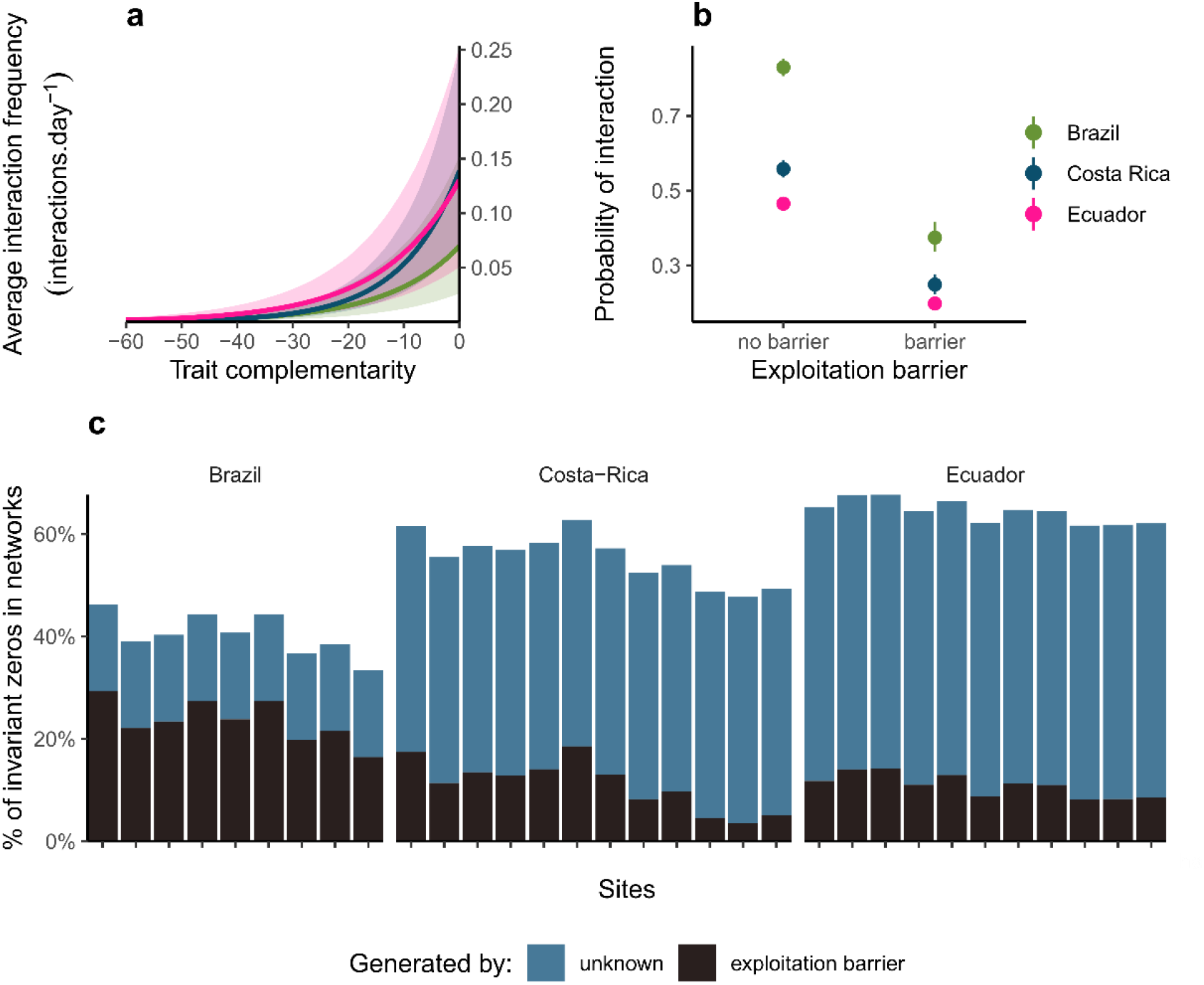
Trait complementarity and exploitation barrier determine plant-hummingbird interactions, and the sparsity of the interaction networks. (a) Predicted average interaction frequencies as a function of trait complementarity between plants and hummingbirds. Ribbons show the 95% confidence interval. (b) Probability of interaction between plants and hummingbirds as a function of the presence of exploitation barrier or not. The probability of interaction relates to the ability of species to interact. For example, in Brazil, a pair of hummingbird and plant species without exploitation barrier between them has 83% of chance to be able to interact together, and 17% of chance to not being able because of unknown mechanism. The chance to be able to interact together drops to 37% if there is an exploitation barrier between them (corolla too long for the bird). Error bars represent 95% confidence interval. (c) Percentage of invariant zeros as a function of the linkage rule generating them: exploitation barrier or unknown process (additional zero-inflation). Within each country, sites are ranked by increasing elevation, from left to right.

In addition to these linkage rules, we found significant zero-inflation (Fig. 2b), highlighted by the fact that even when there was no exploitation barrier between two species, their probability of interacting together was much lower than one. This zero-inflation was captured by an overall intercept, which did not inform about the mechanism, suggesting that we do not know why an important number of pairwise interactions among plants and hummingbirds do not occur. The reason could be imperfect detections, exploitation barriers due to unconsidered traits (e.g. flower orientation and curvature) or any stochastic process (e.g. environmental conditions). Regardless of the mechanism behind this zero-inflation, this showed that interaction networks are highly sparse.

Our model allowed us to distinguish invariant zeros, generated by exploitation barrier and unknown processes, from the “flexible” zeros, generated by trait complementarity and sampling pressure (Fig. S3). On average, 14.5% of possible interactions were forbidden because of exploitation barrier, but with strong variation among sites and countries, from 3.5% in high elevation sites in Costa Rica to 29% in the lowlands of Brazil (Fig. 2c).

Variation in the proportion of forbidden links across sites was explained by the distribution of species traits across sites. We found that flower corolla length tended to decrease with elevation in all countries, although the trend was significantly negative in Costa Rica and Ecuador only (Fig. 3a & S4), while the distribution of hummingbird bill length remained constant over elevation (Fig. 3a & S4; Table S2). This pattern of trait distribution can be translated into an elevation gradient in the proportion of forbidden links: higher elevation communities exhibited fewer forbidden links than lower elevation communities (Fig. 3b & *Table S2)*.

**Figure 3:**
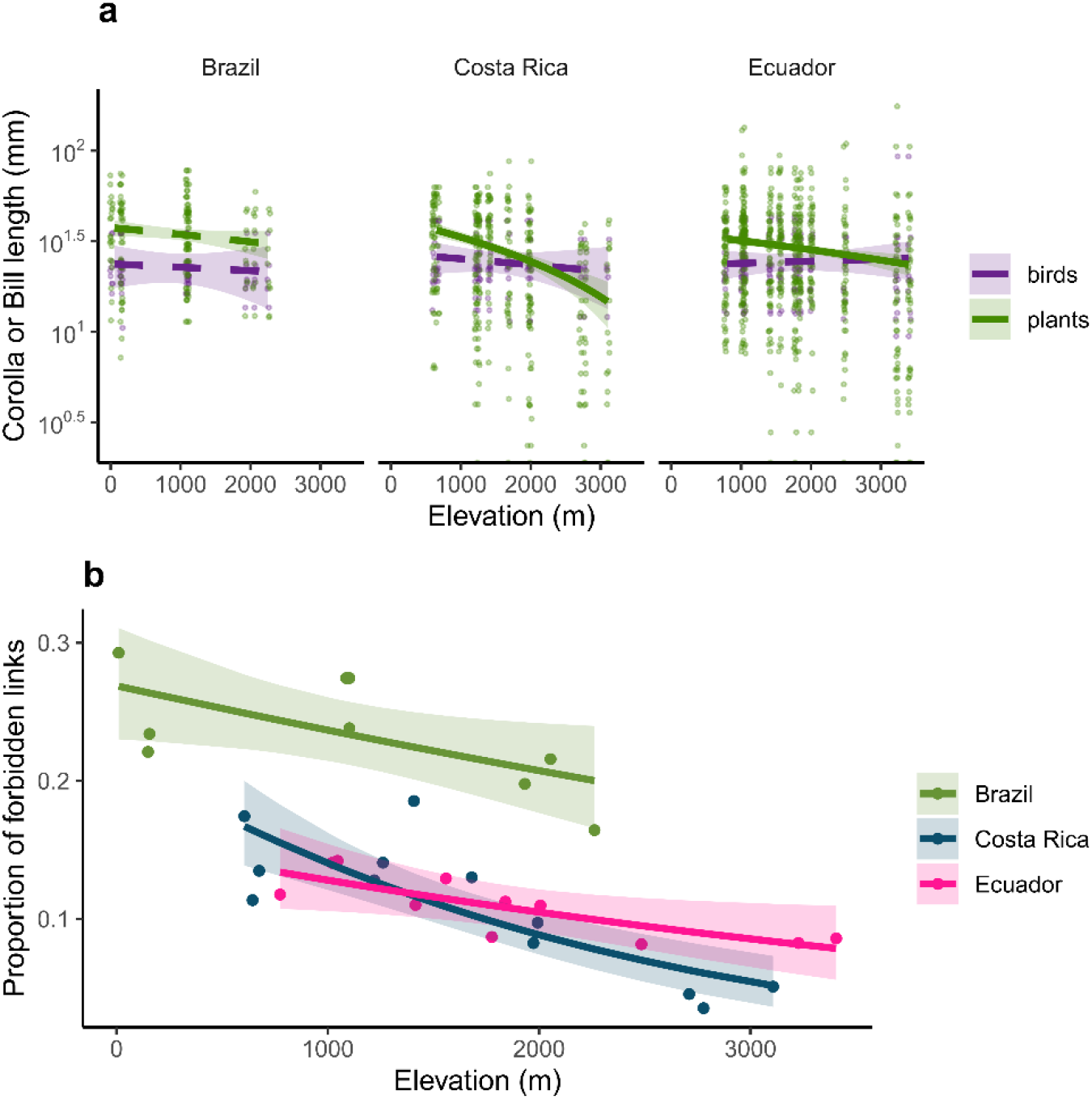
Flower corolla length and proportion of forbidden links decrease over elevation. (a) Corolla or bill length as a function of elevation for every species present at each site, for each country. Note that the y-axis is log-transformed to improve readability. (b) Proportion of forbidden links for each site as a function of elevation and country. In all panels, lines show model predictions and ribbons the associated 95% confidence intervals. Solid lines show trends significantly different from zero while dashed lines show trends non-significantly different zero.

Over the 32 sites studied here, 25 showed a significant decrease of network robustness when accounting for forbidden links, 6 no significant change, and one showed a significant increase of robustness when accounting for forbidden links (Fig. 4 & S5) The higher the proportion of forbidden links in a given site, the more negative the effect on robustness (Fig. 4c). When forbidden links were rare, they tended to increase robustness(Fig. 4c), likely because by preventing the occurrence of some interactions, forbidden links limit the effect of species extinctions to compatible partners.

**Figure 4:**
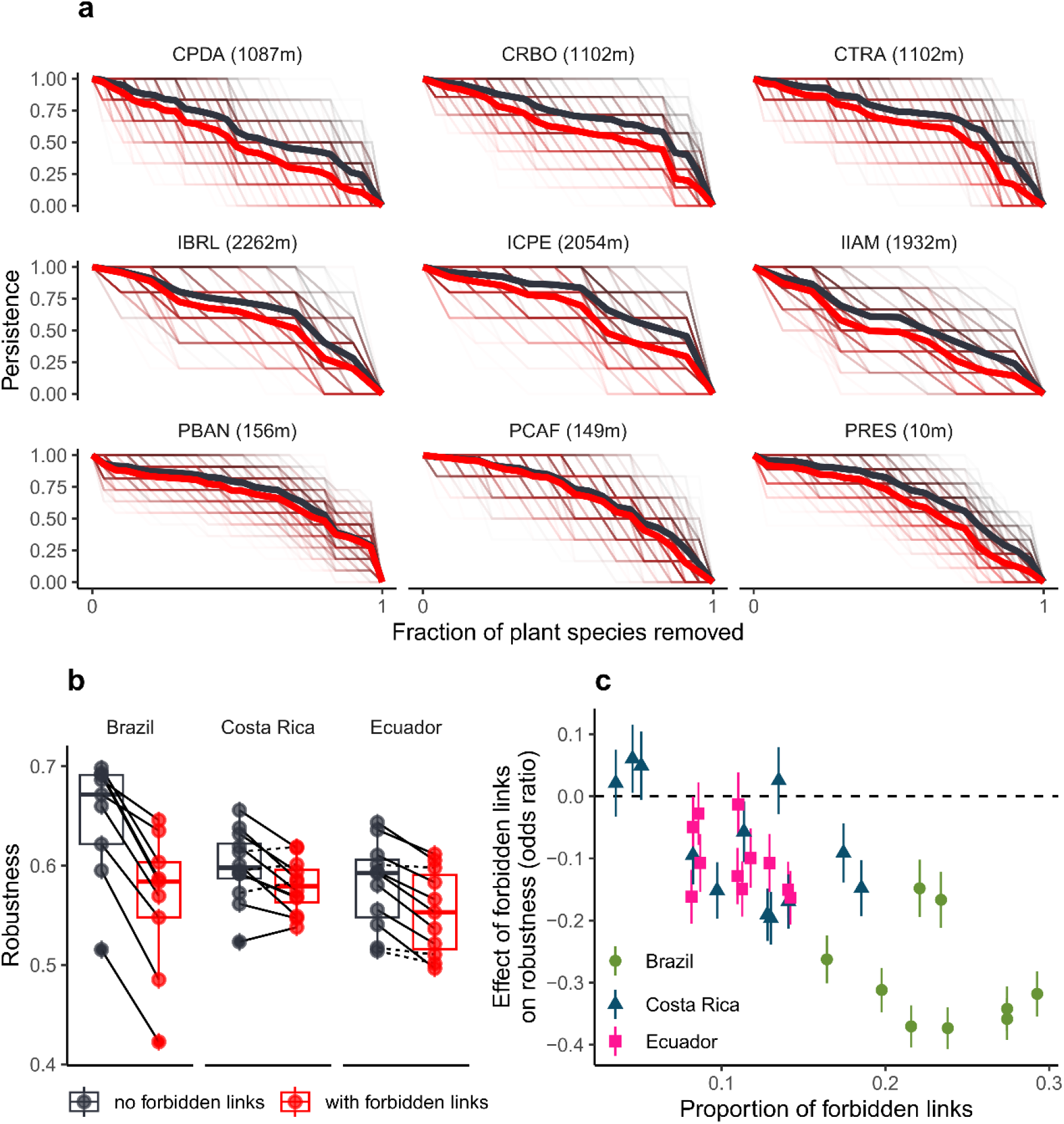
Robustness to plant extinctions is strongly limited by forbidden links, especially at low elevation. (a) Attack tolerance curve for each simulation and each site in Brazil (site acronyms with their elevation, in meters above sea level) when accounting for forbidden links (red) or when neglecting them (black). Thick lines show the average value over the 100 simulations, for each site, while thin lines show individual simulations. (b) Robustness (area under the attack tolerance curve) for each site when accounting for forbidden links (red) or when neglecting them (black). Lines show comparisons within a given site, solid lines show significant change in robustness, dashed lines non-significant change. (c) Effect of forbidden links on robustness (relative to the case without forbidden links), for each site. Error bars are 95% confidence interval. Robustness simulations and values are presented for the scenario “generalists first” and α = 1 (cf. Fig. S5).

This negative effect of forbidden links on network robustness decreased when interaction rewiring rate dropped (Fig. 5). This result is consistent with the fact forbidden links limit robustness to interaction through interaction rewiring and not through modification of the initial structure. Indeed, when rewiring was low (α = 2) forbidden links added few additional constraints on rewiring, and thus had a lower effect on robustness. It is important to note that forbidden links decreases the connectance of initial networks but did not influence their nestedness (Fig. S6).

**Figure 5:**
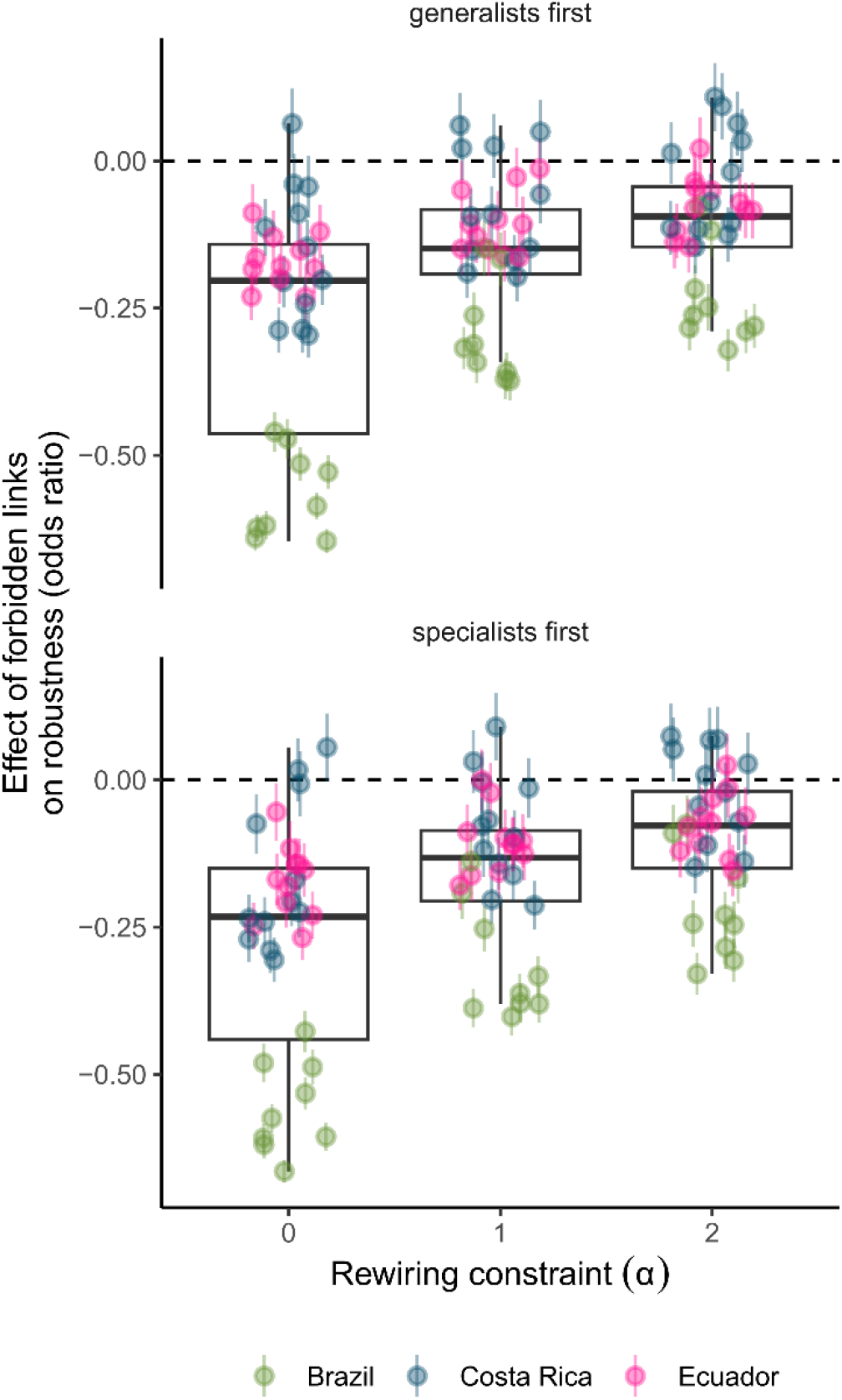
Forbidden links decreased robustness through rewiring. Effect of forbidden links on robustness as a function of rewiring constraint (α), for the two extinction scenarios: either the first plant getting extinct are the more generalists ones, or the more specialist ones. When α = 0 rewiring was not constrained and thus was very high, α = 1 rewiring was moderate, and when α = 2 rewiring was low. Each point is a site and error bars are 95% confidence interval. The boxplots show the overall distributions of effects across sites.

## Discussion

Our results support the idea that forbidden links are not a myth, but instead can be common. Using plant-hummingbird interactions, we complemented existing evidence for forbidden links (Eklöf *et al*. 2013; Jordano *et al*. 2003; Olesen *et al*. 2010; Santamaría & Rodríguez-Gironés 2007) by establishing a statistical and direct link between interactions that have an invariantly low probability of occurring and exploitation barriers. We also show that the prevalence of forbidden links is given by the pool of traits available in the community, leading to considerable variability in their prevalence across environmental gradients (*i*.*e*. elevation). Finally, by using simulations, we found that the higher the proportion of forbidden links, the lower the rewiring in case of local extinction and therefore the lower the network robustness. Since our analyses suggest that exploitation barriers are common in plant-hummingbird interactions, the rescue of species experiencing partner extinction because of interaction rewiring could be much more limited than previously suggested in this system (Sonne *et al*. 2022; Vizentin-Bugoni *et al*. 2020).

We evaluated two aspects of trait matching that are often studied separately, trait complementarity and exploitation barrier, and show that they are both important in determining species interactions. This result is consistent with the hummingbird-plant interaction literature, which have shown that trait complementarity (Dalsgaard *et al*. 2021; Maglianesi *et al*. 2014; Sonne *et al*. 2020) and trait barrier (Vizentin-Bugoni *et al*. 2014) are important factors influencing interaction frequencies, but they have been studied independently. By simultaneously evaluating these two linkage rules, we were able to separate their relative importance, and thus to bring direct statistical evidence for the existence of forbidden links, resulting from exploitation barriers.

The existence of exploitation barriers, producing forbidden links, has been extensively debated (González-Varo & Traveset 2016; Olesen *et al*. 2010). Several studies have shown that they are needed to explain the departure from the scale-free topologies observed in mutualistic networks (Vázquez 2005). However, critiques have been raised about the lack of consideration of spatio-temporal variation as well as intraspecific trait variability that make forbidden links more labile than initially thought, so not really forbidden (González-Varo & Traveset 2016). Here, we partially overcome these limitations by using a probabilistic definition of forbidden links. In our statistical approach, the presence of forbidden links is defined as an invariant decrease in the interaction probability due to exploitation barrier, regardless the context (site, time, etc.) and the sampling pressure. This is important because it means that what we call forbidden links could eventually happen, but if they do, it would be with a much lower probability than an equivalent non-forbidden link. We were able to use a probabilistic approach because our interaction sampling was repeated over different plant individuals from the same species, in different contexts. Thus, even if species traits were used at the species level, interaction data used to estimate exploitation barriers involved intraspecific variation. Our results bridge two points of view: forbidden links are real and potentially important for community structure (Santamaría & Rodríguez-Gironés 2007), but to understand their strength it is important to see them through a probabilistic framework (González-Varo & Traveset 2016).

We also show that the prevalence of exploitation barriers is not constant but varies over space, depending on how species trait distribution changes. We found that, on average, plant corollas are much longer than hummingbird bills at lower elevation, leading to many forbidden links. In the Atlantic forest networks (Brazil), the higher occurrence of forbidden links was mainly due to the fact that understory Bromeliaceae with long-tubed flowers were much more common than in the two other regions. In contrast, at high elevation most of the plant species have relatively short corolla flowers, leading to a decrease in the proportion of forbidden links across elevation. This gradient of flower size over elevation could be due to the fact that large flowers have a large physiological cost at higher elevation, such as water loss (Galen *et al*. 1999). Thus, the cost-benefit balance of producing large flowers could change over elevation (Galen 1999), which can be partially driven by pollinators (Basnett *et al*. 2019; Dalsgaard *et al*. 2009).

Finally, we found that forbidden links strongly decreased the robustness of plant-hummingbird communities to local plant extinctions. Several previous studies have highlighted that robustness to local extinction is determined by the ability of species to rewire interactions lost because of the extinction (Kaiser-Bunbury *et al*. 2010; Ramos-Jiliberto *et al*. 2012; Sheykhali *et al*. 2020; Vizentin-Bugoni *et al*. 2020). Consistent with these results, we also found that network robustness strongly increases as interaction rewiring increases (Fig. S5). However, in the presence of forbidden links rewiring is partially constrained, resulting in a decrease in robustness. Thus, relying on interaction rewiring to buffer the effect of local extinctions due to global change might be a dangerous bet, since rewiring could be more limited than previously expected because of these forbidden links.

It is important to note that we studied forbidden links in terms of mutualistic interactions because we excluded cheating interactions in which hummingbirds pierce the corolla or use an existing hole to rob the nectar. These interactions can allow species to have access to unavailable partners (Duchenne et al. 2023; Rojas-Nossa et al. 2016), and thus blur the limit between forbidden and possible interactions. However, these interactions are likely to be antagonistic rather than mutualistic because plants pay the cost of nectar production but are not pollinated, suggesting that these interactions remain forbidden for mutualism.

Moreover, what we call forbidden links mainly makes sense when focusing on ecological dynamics only. When including evolutionary dynamics, forbidden links would be temporarily forbidden only because species traits can evolve through time. One of the next important steps would be to assess the evolutionary lability of forbidden links and to understand how they shape eco-evolutionary dynamics of natural communities. However, regardless of their evolutionary lability, our study shows that on ecological time scales forbidden links can be common. Assessing the prevalence of these forbidden links across multiple study systems would provide insights on the robustness of ecological communities to the increasing rates of local extinction and colonization (Dornelas *et al*. 2019).

## Acknowledgments

We would like to thank Anna Görlich for her valuable feedback on the manuscript.

In Costa Rica we thank Fabian Monge Badilla, Greivin Serrano Salazar, Samael Padilla, Yandry Hernadez Barboza, Karen Garro Alvarado, Silvia Cascante Chinchilla, Pablo Gutiérrez Campos, Margarita Valverde Quesada, Greilyn Fallas Rodriguez, Lizet Hidalgo Picado, Xiomara Castro Arroyo, Gabriel Zúñiga Morales, Guisselle Arce Soto. We also thank the owners and administrators of the study sites, Los Nimbulos, Los Amigos del Bosque, Villa Mills Experimental Station (ACC), Villa San Miguel, Finca Boquete (Sibu), Bosque del Tolomuco, Centro Biológico Las Quebradas (FUDEBIOL), Las Nubes (York University), Sendero Los Gigantes, Refugio de Aves Los Cusingos (CCT), Hotel Rio Magnolia and Longo Mai.

In Ecuador, data collection was possible thanks to the support of Francisco Tobar, Cristian Poveda and the local assistance of Wilson Hipo, Rolando Hipo, Silvio Calderón, Roberto Pailacho, Willo Vaca, Christian Montalvo, Leo Montalvo, Noé Morales, Edison Tapia, Segundo Imba, Gera Obando, Daniel Ponce. We are also grateful with Jocotoco Foundation, Maldonado family, Antonio Páez, Adela Espinosa, and Alaspungo community. Ministry of Environment in Ecuador provided the research permit Nº 0162019ICFLOFAUDNB/MAE required to conduct field work.

In Brazil, we thank Romulo C. da Silva for help while collecting data. We also thank Mater Natura - Instituto de Estudos Ambientais, especially Helena Zarantonieli and Paulo Pizzi, for administrative support; the Parque Estadual Serra do Mar - Núcleo Picinguaba e Núcleo Cunha - SP, RPPN Instituto Alto Montana - MG, Fazenda Bananal - Paraty - RJ, Parque Nacional do Itatiaia - RJ, for study permit; ICMBIO (research permit 68624-2, 74216-1, 69412-1) and the Instituto de Pesquisas Ambientais (research permit SMA N. 260108 – 005.133/2019) and SISGEN (access registration A81367A); CAPES (Finance Code 001 to ARG, BGR, DSV, LLR, RO and DB), CNPQ (grant CAPES-PRINT - 88887.756715/2022-00, and 312580/2020- 7 to IGV).

The project was funded by the European Research Council (ERC) under the European Union’s Horizon 2020 research and innovation program (grant agreement N° 787638) and by the Swiss National Science Foundation (grant Nº 173342).

## References

Albert, R., Jeong, H. & Barabási, A.-L. (2000). Error and attack tolerance of complex networks. Nature, 406, 378–382.

Bartomeus, I., Gravel, D., Tylianakis, J.M., Aizen, M.A., Dickie, I.A. & Bernard-Verdier, M. (2016). A common framework for identifying linkage rules across different types of interactions. Funct. Ecol., 30, 1894–1903.

Bascompte, J. (2009). Disentangling the Web of Life. Science, 325, 416–419.

Basnett, S., Ganesan, R. & Devy, S.M. (2019). Floral traits determine pollinator visitation in Rhododendron species across an elevation gradient in the Sikkim Himalaya. Alp. Bot., 129, 81–94.

Busiello, D.M., Suweis, S., Hidalgo, J. & Maritan, A. (2017). Explorability and the origin of network sparsity in living systems. Sci. Rep., 7, 12323.

CaraDonna, P.J., Petry, W.K., Brennan, R.M., Cunningham, J.L., Bronstein, J.L., Waser, N.M., et al. (2017). Interaction rewiring and the rapid turnover of plant–pollinator networks. Ecol. Lett., 20, 385–394.

Dalsgaard, B., Martín González, A.M., Olesen, J.M., Ollerton, J., Timmermann, A., Andersen, L.H., et al. (2009). Plant–hummingbird interactions in the West Indies: floral specialisation gradients associated with environment and hummingbird size. Oecologia, 159, 757–766.

Dalsgaard, B., Maruyama, P.K., Sonne, J., Hansen, K., Zanata, T.B., Abrahamczyk, S., et al. (2021). The influence of biogeographical and evolutionary histories on morphological trait-matching and resource specialization in mutualistic hummingbird–plant networks. Funct. Ecol., 35, 1120–1133.

Doré, M., Fontaine, C. & Thébault, E. (2021). Relative effects of anthropogenic pressures, climate, and sampling design on the structure of pollination networks at the global scale. Glob. Change Biol., 27, 1266–1280.

Dornelas, M., Gotelli, N.J., Shimadzu, H., Moyes, F., Magurran, A.E. & McGill, B.J. (2019). A balance of winners and losers in the Anthropocene. Ecol. Lett., 22, 847–854.

Duchenne, F., Aubert, S., Barreto, E., Brenes, E., Maglianesi, M.A., Santander, T., et al. (2023). When cheating turns into a stabilizing mechanism of mutualistic networks.

Eklöf, A., Jacob, U., Kopp, J., Bosch, J., Castro-Urgal, R., Chacoff, N.P., et al. (2013). The dimensionality of ecological networks. Ecol. Lett., 16, 577–583.

Galen, C. (1999). Why Do Flowers Vary?: The functional ecology of variation in flower size and form within natural plant populations. BioScience, 49, 631–640.

Galen, C., Sherry, R.A. & Carroll, A.B. (1999). Are flowers physiological sinks or faucets? Costs and correlates of water use by flowers of Polemonium viscosum. Oecologia, 118, 461–470.

Gonzalez, O. & Loiselle, B.A. (2016). Species interactions in an Andean bird–flowering plant network: phenology is more important than abundance or morphology. PeerJ, 4, e2789.

González-Varo, J.P. & Traveset, A. (2016). The Labile Limits of Forbidden Interactions. Trends Ecol. Evol., 31, 700–710.

Gravel, D., Poisot, T., Albouy, C., Velez, L. & Mouillot, D. (2013). Inferring food web structure from predator–prey body size relationships. Methods Ecol. Evol., 4, 1083–1090.

Jordano, P. (2016). Sampling networks of ecological interactions. Funct. Ecol., 30, 1883–1893.

Jordano, P., Bascompte, J. & Olesen, J.M. (2003). Invariant properties in coevolutionary networks of plant–animal interactions. Ecol. Lett., 6, 69–81.

Junker, R.R., Blüthgen, N., Brehm, T., Binkenstein, J., Paulus, J., Schaefer, H.M., et al. (2013). Specialization on traits as basis for the niche-breadth of flower visitors and as structuring mechanism of ecological networks. Funct. Ecol., 27, 329–341.

Kaiser-Bunbury, C.N., Muff, S., Memmott, J., Müller, C.B. & Caflisch, A. (2010). The robustness of pollination networks to the loss of species and interactions: a quantitative approach incorporating pollinator behaviour. Ecol. Lett., 13, 442–452.

Kéfi, S., Miele, V., Wieters, E.A., Navarrete, S.A. & Berlow, E.L. (2016). How Structured Is the Entangled Bank? The Surprisingly Simple Organization of Multiplex Ecological Networks Leads to Increased Persistence and Resilience. PLOS Biol., 14, e1002527.

Klumpers, S.G.T., Stang, M. & Klinkhamer, P.G.L. (2019). Foraging efficiency and size matching in a plant–pollinator community: the importance of sugar content and tongue length. Ecol. Lett., 22, 469–479.

Machado-de-Souza, T., Campos, R.P., Devoto, M. & Varassin, I.G. (2019). Local drivers of the structure of a tropical bird-seed dispersal network. Oecologia, 189, 421–433.

Maglianesi, M.A., Blüthgen, N., Böhning-Gaese, K. & Schleuning, M. (2014). Morphological traits determine specialization and resource use in plant–hummingbird networks in the neotropics. Ecology, 95, 3325–3334.

McGuire, J.A., Witt, C.C., Remsen, J.V., Corl, A., Rabosky, D.L., Altshuler, D.L., et al. (2014). Molecular Phylogenetics and the Diversification of Hummingbirds. Curr. Biol., 24, 910–916.

Memmott, J., Waser, N.M. & Price, M.V. (2004). Tolerance of pollination networks to species extinctions. Proc. R. Soc. Lond. B Biol. Sci., 271, 2605–2611.

Olesen, J.M., Bascompte, J., Dupont, Y.L., Elberling, H., Rasmussen, C. & Jordano, P. (2010). Missing and forbidden links in mutualistic networks. Proc. R. Soc. B Biol. Sci., 278, 725–732.

Paton, D.C. & Collins, B.G. (1989). Bills and tongues of nectar-feeding birds: A review of morphology, function and performance, with intercontinental comparisons. Aust. J. Ecol., 14, 473–506.

Paudel, B.R., Shrestha, M., Burd, M., Adhikari, S.Sun, Y.-S. & Li, Q.-J. (2016). Coevolutionary elaboration of pollination-related traits in an alpine ginger (Roscoea purpurea) and a tabanid fly in the Nepalese Himalayas. New Phytol., 211, 1402–1411.

Pocock, M.J.O., Evans, D.M. & Memmott, J. (2012). The Robustness and Restoration of a Network of Ecological Networks. Science, 335, 973–977.

Ramos-Jiliberto, R., Valdovinos, F.S., Moisset de Espanés, P. & Flores, J.D. (2012). Topological plasticity increases robustness of mutualistic networks. J. Anim. Ecol., 81, 896–904.

Rojas-Nossa, S.V., Sánchez, J.M. & Navarro, L. (2016). Nectar robbing: a common phenomenon mainly determined by accessibility constraints, nectar volume and density of energy rewards. Oikos, 125, 1044–1055.

Santamaría, L. & Rodríguez-Gironés, M.A. (2007). Linkage Rules for Plant–Pollinator Networks: Trait Complementarity or Exploitation Barriers? PLOS Biol., 5, e31.

Sazatornil, F.D., Moré, M., Benitez-Vieyra, S., Cocucci, A.A., Kitching, I.J., Schlumpberger, B.O., et al. (2016). Beyond neutral and forbidden links: morphological matches and the assembly of mutualistic hawkmoth–plant networks. J. Anim. Ecol., 85, 1586–1594.

Sheykhali, S., Fernández-Gracia, J., Traveset, A., Ziegler, M., Voolstra, C.R., Duarte, C.M., et al. (2020). Robustness to extinction and plasticity derived from mutualistic bipartite ecological networks. Sci. Rep., 10, 9783.

Song, C., Von Ahn, S., Rohr, R.P. & Saavedra, S. (2020). Towards a Probabilistic Understanding About the Context-Dependency of Species Interactions. Trends Ecol. Evol., 35, 384–396.

Sonne, J., Maruyama, P.K., Martín González, A.M., Rahbek, C., Bascompte, J. & Dalsgaard, B. (2022). Extinction, coextinction and colonization dynamics in plant–hummingbird networks under climate change. Nat. Ecol. Evol., 6, 720–729.

Sonne, J., Vizentin-Bugoni, J., Maruyama, P.K., Araujo, A.C., Chávez-González, E., Coelho, A.G., et al. (2020). Ecological mechanisms explaining interactions within plant– hummingbird networks: morphological matching increases towards lower latitudes. Proc. R. Soc. B Biol. Sci., 287, 20192873.

Stang, M., Klinkhamer, P.G.L., Waser, N.M., Stang, I. & van der Meijden, E. (2009). Size-specific interaction patterns and size matching in a plant–pollinator interaction web. Ann. Bot., 103, 1459–1469.

Su, Y. & Yajima, M. (2012). Package ‘R2jags’. A Package for Running jags from R.

Thébault, E. & Fontaine, C. (2010). Stability of Ecological Communities and the Architecture of Mutualistic and Trophic Networks. Science, 329, 853–856.

Tylianakis, J.M. & Morris, R.J. (2017). Ecological Networks Across Environmental Gradients. Annu. Rev. Ecol. Evol. Syst., 48, 25–48.

Vázquez, D.P. (2005). Degree distribution in plant–animal mutualistic networks: forbidden links or random interactions? Oikos, 108, 421–426.

Vázquez, D.P., Blüthgen, N., Cagnolo, L. & Chacoff, N.P. (2009a). Uniting pattern and process in plant–animal mutualistic networks: a review. Ann. Bot., 103, 1445–1457.

Vázquez, D.P., Chacoff, N.P. & Cagnolo, L. (2009b). Evaluating multiple determinants of the structure of plant–animal mutualistic networks. Ecology, 90, 2039–2046.

Vizentin-Bugoni, J., Debastiani, V.J., Bastazini, V.A.G., Maruyama, P.K. & Sperry, J.H. (2020). Including rewiring in the estimation of the robustness of mutualistic networks. Methods Ecol. Evol., 11, 106–116.

Vizentin-Bugoni, J., Maruyama, P.K. & Sazima, M. (2014). Processes entangling interactions in communities: forbidden links are more important than abundance in a hummingbird– plant network. Proc. R. Soc. B Biol. Sci., 281, 20132397.

Week, B. & Nuismer, S.L. (2021). Coevolutionary Arms Races and the Conditions for the Maintenance of Mutualism. Am. Nat., 198, 195–205.

Weinstein, B.G. (2018). Scene-specific convolutional neural networks for video-based biodiversity detection. Methods Ecol. Evol., 9, 1435–1441.

Zhao, Y.-H., Lázaro, A., Li, H.-D., Tao, Z.-B., Liang, H., Zhou, W., et al. (2022). Morphological trait-matching in plant–Hymenoptera and plant–Diptera mutualisms across an elevational gradient. J. Anim. Ecol., 91, 196–209.

