## Supplementary Materials for "A probabilistic view of forbidden links: their prevalence and their consequences for the robustness of plant-hummingbird communities"

F. Duchenne *et al.*

### Supplementary Methods

*Morphological traits of plants and hummingbirds*

Hummingbird bill lengths (i.e. length of exposed culmen) were compiled from literature (Bueno *et al.* 2021; Dalsgaard *et al.* 2021; Graham *et al.* 2012) and measured on the field. Bill length measures were averaged to get one value per species, with the number of individual measures varying from 1 to 160 (median = 8). For one hummingbird species (*Chlorostilbon russatus*), the length of exposed culmen (distance from the bill tip to the point where the tips of the forehead feathers begin to hide the culmen) was inferred from the length of the bill tip from the notch on the forehead. To do so, we used a linear model explaining all available measures of the length of exposed culmen by the length of the bill tip from the notch on the forehead. This model explained well the length of exposed culmen ($R^2 = 0.94$), thus we used it to predict the missing value of exposed culmen.

Plant corolla lengths were measured on pictures with scale from plants collected in the field using Image J (Schneider et al. 2012). Corolla lengths were averaged to get one value per species, with the number of individual measures varying from 1 to 46 (median = 5).

*Census of the present flowers*

In each site all flowering plant species with an ornithophilous syndrome, as well as those on which either observers or cameras detected foraging hummingbirds, were recorded for each month along the 1.5 km transect at each site. All flowers within five meters on either side of the transect were counted. Thus, we obtained a list of flowering plant species present for each month and site.

*Accounting for phenologies and abundances when estimating linkage rules*

Our sampling events were spread over time; thus, we needed species phenologies to account for the temporal variation in interactions due to the seasonal turnover of species. Since we sampled open flowers only, we do not need to account for flowering phenologies. To account for hummingbirds' phenology, for each hummingbird species and month, we calculated the fraction of interactions that a given month represents relative to the maximum number of interactions detected on the focal site during a given month. This gave us an index of relative abundance, for each hummingbird species *i*, each site *s*, and each month *m*. We assumed that hummingbirds' lifespan is longer than a year, so the phenological variation in their abundances among months was due to spatial movements rather than to a birth-death process.

*Priors for the Bayesian model*

We assumed that zeros can be over-represented in the distribution of interaction frequencies (zero-inflated) because of stochastics process or exploitation barriers due to missing traits, so we let the value of $\beta_0$ varying. We used $\mathcal{N}(0,2)$ for $\beta_0$ and $\beta_1$ following the recommendations for "uninformative" priors in occupancy models (Northrup & Gerber 2018).

The prior distribution for fixed effects of the equation (2), $\beta_{2...4}$, was $\mathcal{N}(0,10)$. For random effects, we used half-Cauchy hyperpriors (Student's t-distribution with 1 degree of freedom) to model the standard deviations. For $\rho$, the scale parameter of spatial correlation, we used a gamma distribution with the shape parameter of 3 and a rate parameter of 0.1. For the overdispersion parameter, we used a Negative binomial distribution, $r \sim \text{NB}(4{,}0.2)$.

*Percentage of zeros generated by each linkage rule*

The percentage of zeros generated by the unknown process (additional zero-inflation) was evaluated for each site as $\frac{100}{n_h n_p} \sum_{i=1}^{n_h} \sum_{j=1}^{n_p} (1 - \frac{e^{\beta_0}}{1+e^{\beta_0}})$, where $n_p$ and $n_h$ were the number of plant and hummingbird species, respectively, present on that site.

For each site, the percentage of zeros generated by exploitation barrier was evaluated as $\frac{100}{n_h n_p} \sum_{i=1}^{n_h} \sum_{j=1}^{n_p} \frac{e^{\beta_0}}{1+e^{\beta_0}} - \psi_{ji}$, and the percentage of zeros generated by trait complementarity as $\frac{100}{n_h n_p} \sum_{i=1}^{n_h} \sum_{j=1}^{n_p} \psi_{ji} \times \text{P}[\text{X} = 0]$, where $\text{X} \sim \text{NB}(r, p_{jis})$, the negative binomial distribution from the Bayesian model, equation (2). To get an estimate that was aggregated over seasons, we used a value of $p_{jis}$ for a value of phenological match averaged over the 12 months of the year, for each pair of plant-hummingbird species ($\overline{\text{pheno}_{jis}} = \frac{1}{12} \sum_{m=1}^{12} \text{pheno}_{js} \times \text{pheno}_{is}$). Since $\lambda_{jis}$ also depends on sampling duration, we estimated the percentage of zeros generated by each linkage rule as a function of sampling duration.

*Predict plant-hummingbird interaction networks for each site*

To assess robustness, we wanted to start simulations of extinctions on initial networks that did not depend on sampling pressure. We used our statistical model to predict the average interaction count over 12 hours of daylight, which is roughly equal to a day in the tropics. Also, the way we measured robustness does not require us to parametrize complex dynamic models and associated time-consuming simulations. For simplicity, this approach models species as present (persisting) or absent (extinct) but neglects more complex population dynamics and variations. To make initial interaction networks consistent with our approach, we used our model to predict average initial interaction networks aggregated over time.

Our model gave average estimates of interaction per unit of time per camera (= one flower or inflorescence), $\lambda_{jismvc}$ which depends on time (month *m* and visit *v*) on the sampling pressure (duration of sampling of camera *c*). To focus on the underlying interaction matrix corrected by abundances and time, we measured interaction strength independent from time effects and from hummingbird abundances (model absorbed through $\theta_{si}$). To do so, while predicting we neglected the bird-site and temporal random effects. Additionally, we wanted to obtain an interaction network aggregated over seasons, so we predicted interaction frequencies for a value of phenological match averaged over the 12 months, for each pair of plant-hummingbird species ($\overline{\text{pheno}_{jis}} = \frac{1}{12} \sum_{m=1}^{12} \text{pheno}_{js} \times \text{pheno}_{is}$). We kept the plant and site random effects that captured mostly variation in terms of attractivity for hummingbirds of plant species and sites.

We used the following equation from the “NB model” to predict interaction frequencies:

**Exploitation barrier submodel**

$$\psi_{ji} \, ; logit(\psi_{ji}) = \beta_0 + \beta_1 \times barrier_{ij} \qquad (S1)$$

**Trait complementarity submodel**

$$\lambda_{jis} ;$$

$$log(\lambda_{jis}) = \beta_2 + \beta_3 \times compl._{ij} + \beta_4 \times \overline{\text{pheno}_{\jmath\imath s}} + log\ (12) + \theta_j + \theta_s \qquad (S2)$$

**Prediction**

$$\psi_{ji}\lambda_{jis} \qquad (S3)$$

When a plant was present on a site but was absent from the model because its interactions had not been recorded with camera traps, we predicted interaction frequencies while ignoring the plant species' random effect ($\theta_j = 0$). As a final measure of interaction strength, we used $\psi_{ji}\lambda_{jis}$. When we wanted to ignore forbidden links in simulations to assess robustness, we used $logit(\psi_{ji}) = \beta_0$ instead of $logit(\psi_{ji}) = \beta_0 + \beta_1 \times barrier_{ij}$ in equation (S1).

*Network robustness simulations*

First, we removed a plant species (*j*), starting with the most generalist one (highest degree), and all its interactions from the network to simulate extinction. Second, we allowed all hummingbird species (*i*) that lost interactions because of the extinction of plant *j* to rewire a percentage ($R_i$) of these lost interactions, according to the following formula:

$$R_i = \frac{1}{n_p}\sum_{k=1}^{n_p} \psi_{ji} e^{-\alpha \frac{\sqrt{(compl._{ij} - compl._{ik})^2}}{d_{max}}} \qquad (S4)$$

Equation (S4) says that the number of interactions lost for hummingbird *i* because of extinction of plant *j* that was rewired was proportional to the arithmetic mean of the pairwise similarities between plant *j* and other plant species, in terms of trait complementarity with hummingbird *j*, weighted by their probability of interaction ($\psi_{ji}$, from equation (1)). In other words, the more different plant *j* was, in terms of complementarity with hummingbird *i*, the less able the focal hummingbird to rewire the interaction lost because of the extinction of plant *j*. As shown in equation (5), the similarity between two plants decreased exponentially by a rate $\alpha$ ($\alpha \geq 0$) with the Euclidean distance between their complementarity values with hummingbird *i*, normalized by the higher maximum distance ($d_{max}$). The lower $\alpha$ was, the easier the rewiring was. By normalizing complementarity distances by $d_{max}$, we erased differences among sites in terms of absolute values of trait complementarity.

Third, for each hummingbird species, we drew its survival status from a Bernoulli distribution, parametrized with a probability of survival equal to one minus the probability of extinction ($p_{ext_i}$), calculated with the following formula:

$$p_{ext_i} = \frac{(1 - R_i) I_{ji}}{\sum_{k=1}^{n_p} I_{ki}} \qquad (S5)$$

A hummingbird had a probability of extinction proportional to the fraction of interactions it lost relative to the initial state. After possible secondary extinctions, the second most generalist plant species was removed, and secondary extinctions were simulated as before, and so on until the last plant was removed. The curve of the fraction of hummingbird species persisting against the fraction of plant species removed is the attack tolerance curve. The area under this curve is the measure of robustness. It is bounded between 0, the lowest robustness possible, and 1, the highest possible. We estimated the robustness for each site according to the area under the attack tolerance curve for the 500 simulations.

We performed these 500 simulations for two cases, with ($logit(\psi_{ji}) = \beta_0 + \beta_1 \times barrier_{ij}$) or without ($logit(\psi_{ji}) = \beta_0$) forbidden links. To see how our results depended on the sequence of extinctions and strength of rewiring, we also performed these simulations for two different scenarios of plant extinctions, removing plant species from the more generalists (highest degree) to the more specialists' one (lowest degree) or the opposite, and for three different values of $\alpha$ (0, 1 and 2).

*Network metrics*

To see if initial network structure changed with the inclusion of forbidden links, we measured the connectance and nestedness. Connectance was measured as the sum of links divided by the number of possible interactions. Nestedness was measured using the Weighted-Interaction Nestedness Estimator (Galeano et al. 2009) usingthe "bipartite" R packages (Dormann *et al.* 2008)

*References*

Bueno, R. de O., Zanata, T.B. & Varassin, I.G. (2021). Niche partitioning between hummingbirds and well-matched flowers is independent of hummingbird traits. *Journal of Tropical Ecology*, 37, 193–199.

Dalsgaard, B., Maruyama, P.K., Sonne, J., Hansen, K., Zanata, T.B., Abrahamczyk, S., *et al.* (2021). The influence of biogeographical and evolutionary histories on morphological trait-matching and resource specialization in mutualistic hummingbird–plant networks. *Functional Ecology*, 35, 1120–1133.

Dormann, C.F., Gruber, B. & Fründ, J. (2008). Introducing the bipartite Package: Analysing Ecological Networks. *R News*, 8, 4.

Galeano, J., Pastor, J.M. & Iriondo, J.M. (2009). Weighted-Interaction Nestedness Estimator (WINE): A new estimator to calculate over frequency matrices. *Environmental Modelling & Software*, 24, 1342–1346.

Graham, C.H., Parra, J.L., Tinoco, B.A., Stiles, F.G. & McGuire, J.A. (2012). Untangling the influence of ecological and evolutionary factors on trait variation across hummingbird assemblages. *Ecology*, 93, S99–S111.

Northrup, J.M. & Gerber, B.D. (2018). A comment on priors for Bayesian occupancy models. *PLOS ONE*, 13, e0192819.

Schneider, C.A., Rasband, W.S. & Eliceiri, K.W. (2012). NIH Image to ImageJ: 25 years of image analysis. *Nat Methods*, 9, 671–675.

### Supplementary Figures

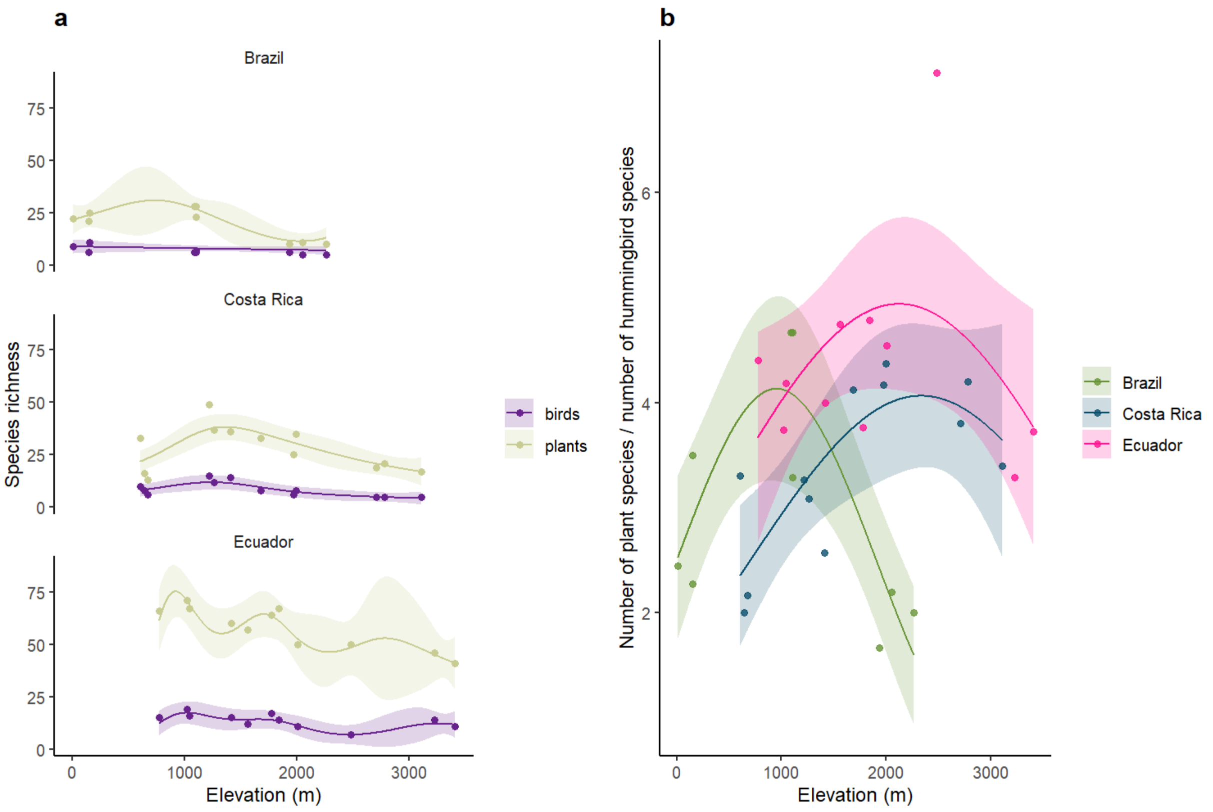

***Figure S1: Species richness and network asymmetry over elevation.*** *(a) Species richness for hummingbirds (purple, "birds") and plants (green) as a function of elevation and country. Lines and ribbons show predictions and associated 95% confidence intervals of a generalized additive model with a Poisson error structure. (b) Network asymmetry (number of plant species divided by number of hummingbird species present at each site) as a function of elevation (in meters above sea level) and country.*

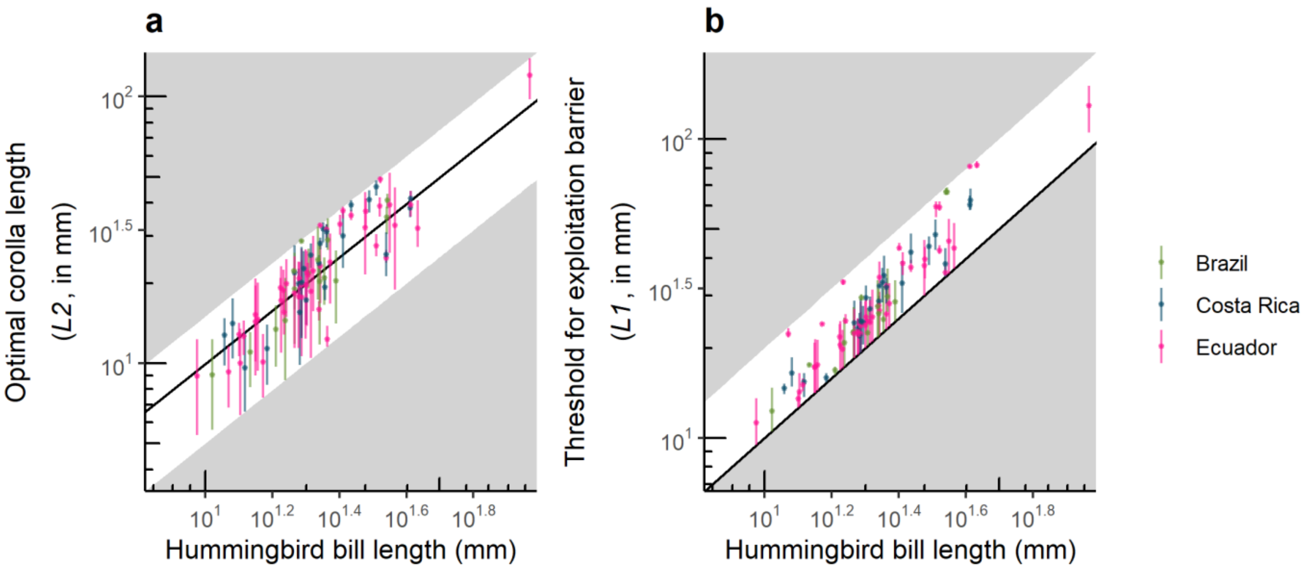

***Figure S2: Values of the latent variables used to model trait complementarity and exploitation barrier.*** *Predictions of (a) the optimal corolla length for trait complementarity and (b) the maximal corolla length (above which exploitation barrier occurs) as a function of bill length, for each hummingbird species. The white areas represent the area of the possible values for L2 and L1, respectively. The black solid line is the first bisector. In all panels, ribbons and error bars represent 95% confidence interval associated with predictions. Note that axis are log-transformed to represent the hummingbird species with a very long bill (Sword-billed hummingbird) on the same plot as the other species. The colors corresponds to country: pink = Ecuador, blue = Costa Rica and green = Brazil. Each point is a hummingbird species and country combination. In our dataset, Costa Rica and Ecuador shared 9 common species, while they each did not share any hummingbird species in common with Brazil.*

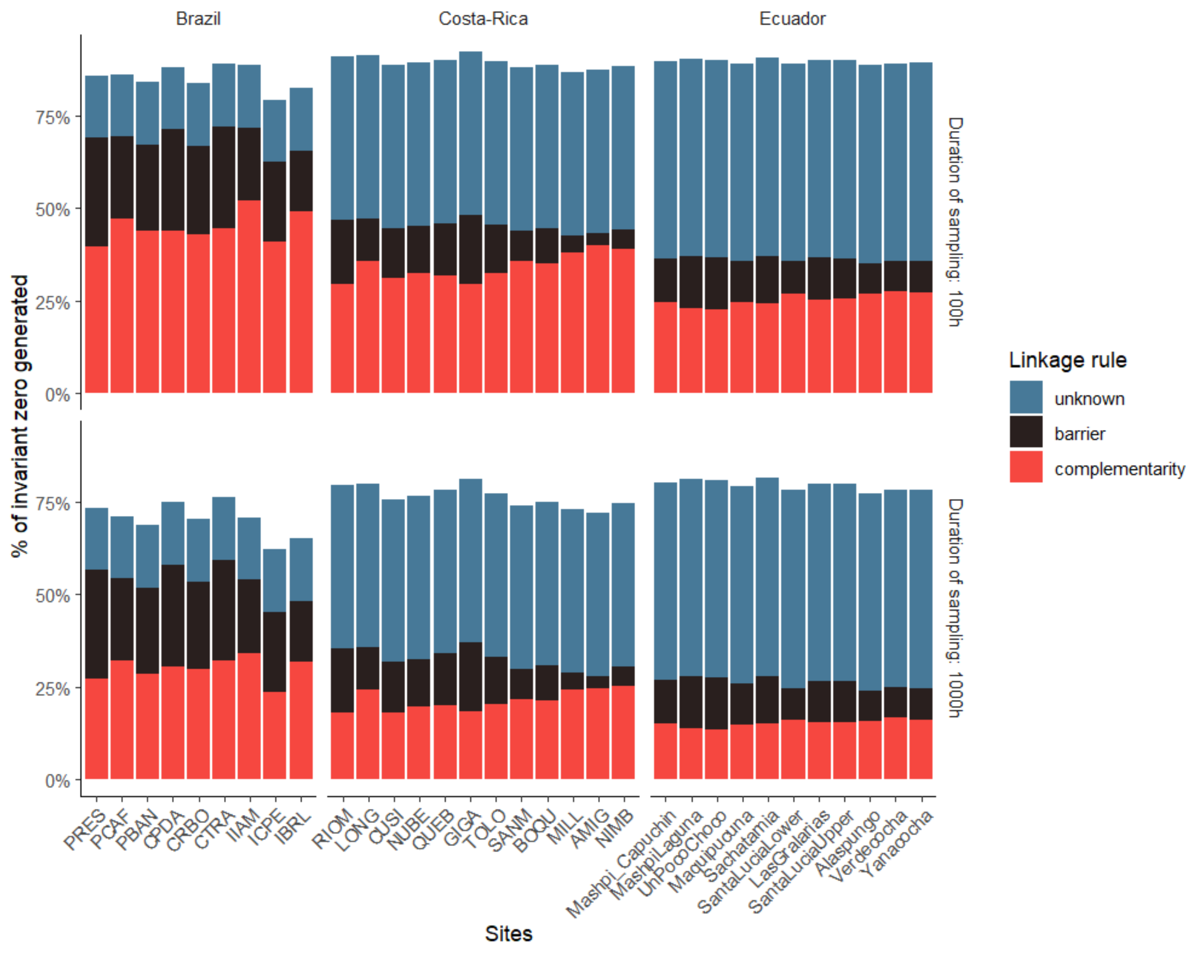

***Figure S3: The sparsity of ecological networks and their causes.*** *The proportion of zero in interaction frequencies generated by each linkage rule for each site. Within each country, sites are ranked by increasing elevation, from left to right. The first row shows the proportion of zero generated by each linkage rule when predicting data with a sampling duration of 100h for each camera, and the second row when predicting data with a sampling duration of 1000h for each camera. As trait complementarity is a continuous mechanism, the proportion of zero it generates tends to zero when sampling pressure increases.*

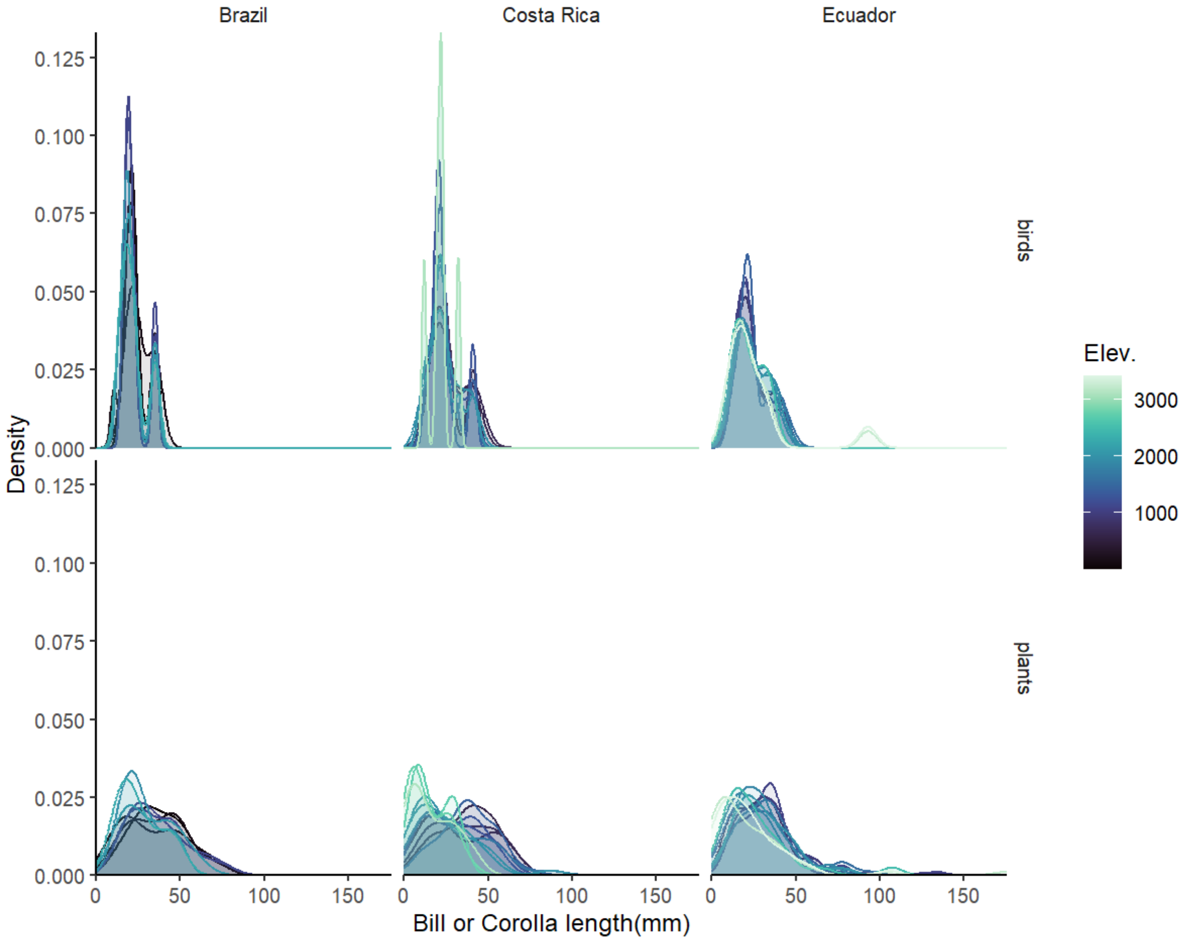

***Figure S4: Trait distributions over elevation and countries.*** *Distribution of species trait values for each site, as a function of elevation (Elev., in meters above sea level) and country.*

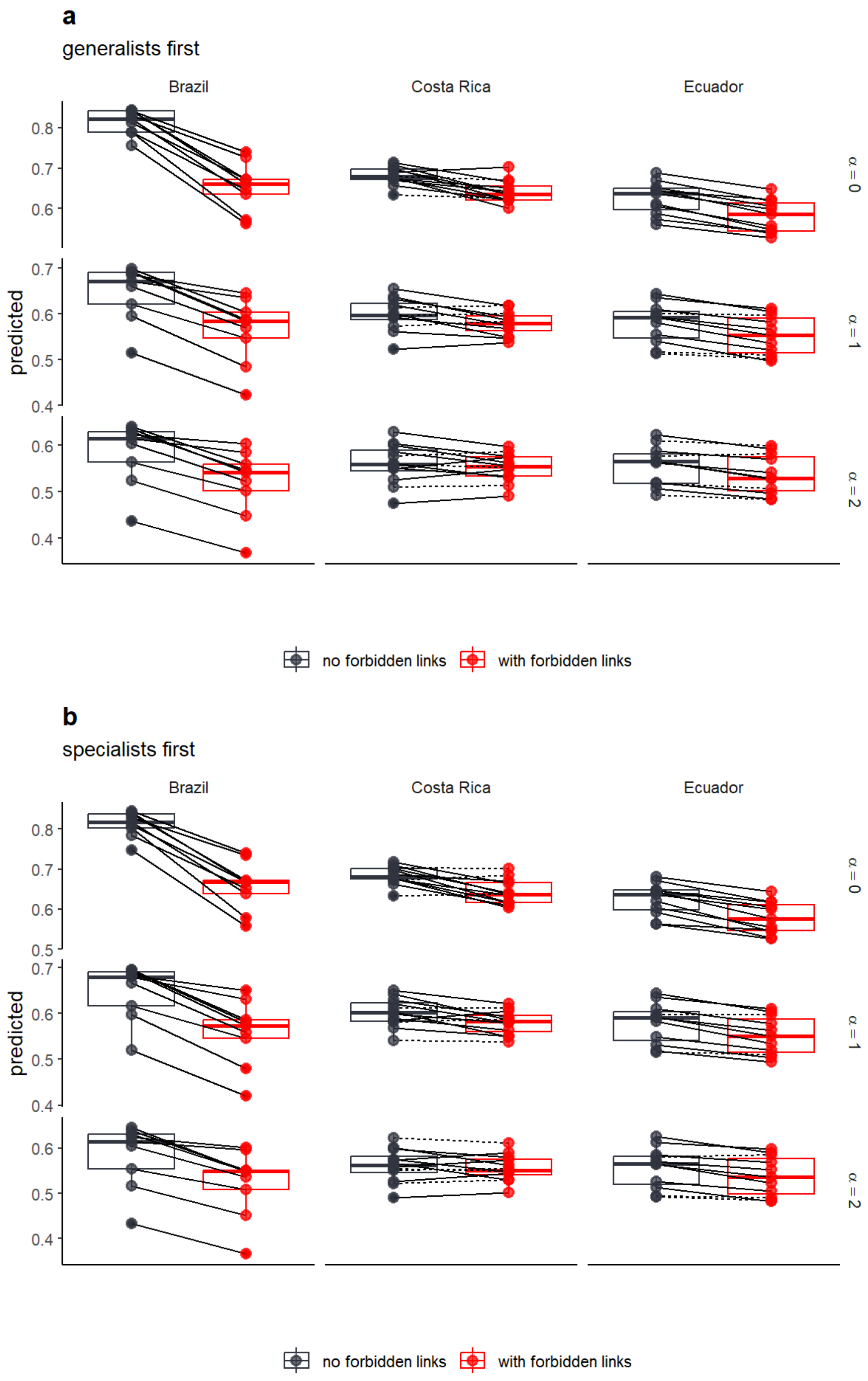

***Figure S5: Sensitivity analyses of robustness simulations.*** *Same figure as Fig. 4b of the main text but for different values of interaction rewiring, from α = 0 (complete rewiring if no zero-inflation) to α = 2 (rewiring strongly constrained by trait matching), and for a different scenario, (a) when removing the most generalist plant first or (b) when removing the most specialist plant first. Robustness (area under the attack tolerance curve) when accounting for forbidden links (red) or when neglecting them (dark grey). Lines show comparisons within a given site, solid lines show significant change in robustness, dashed lines non-significant change.*

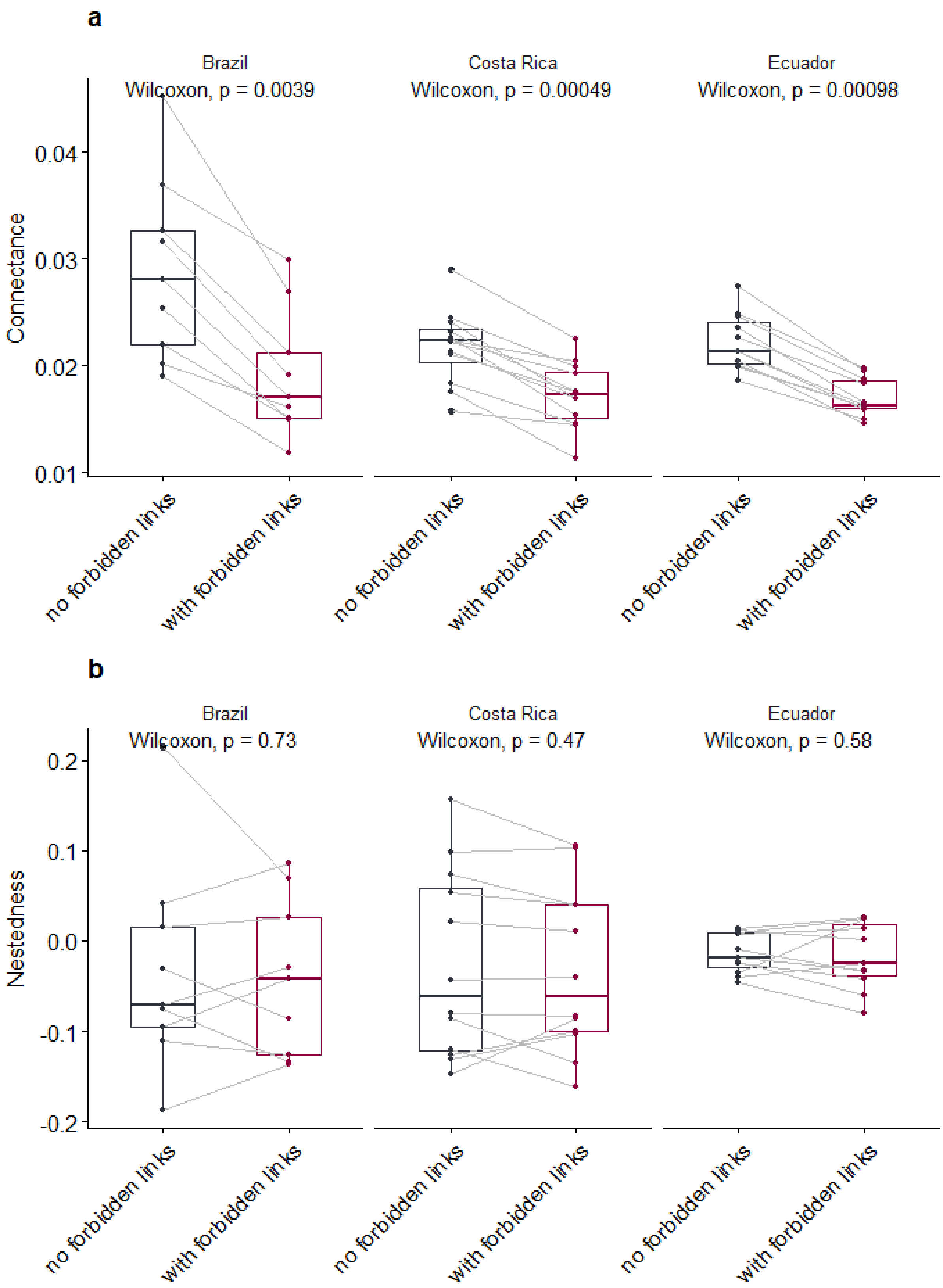

***Figure S6: Changes in network structure when including forbidden links, for each country.*** *Network metric, (a) connectance and (b) nestedness, as a function of the presence of forbidden links or not. Each point is a site, and grey lines link identical sites, with or without forbidden links. The p-value is the p-value from a paired-sample Wilcoxon test.*

### Supplementary Tables

Table S1: Location and elevation of the studied sites. Elevation is in meters above sea level.

| Site | Longitude | Latitude | Elevation | Country |
|---|---|---|---|---|
| **PRES** | -44.8344 | -23.3651 | 10 | Brazil |
| **PCAF** | -44.8258 | -23.3268 | 149 | Brazil |
| **PBAN** | -44.775 | -23.2108 | 156 | Brazil |
| **CPDA** | -45.0246 | -23.2509 | 1087 | Brazil |
| **CRBO** | -45.0138 | -23.2446 | 1102 | Brazil |
| **CTRA** | -45.0232 | -23.2342 | 1102 | Brazil |
| **IIAM** | -44.815 | -22.3767 | 1932 | Brazil |
| **ICPE** | -44.7392 | -22.3665 | 2054 | Brazil |
| **IBRL** | -44.7325 | -22.3581 | 2262 | Brazil |
| **RIOM** | -83.7821 | 9.340159 | 605 | Costa Rica |
| **LONG** | -83.4882 | 9.257907 | 644 | Costa Rica |
| **CUSI** | -83.6281 | 9.32995 | 675 | Costa Rica |
| **NUBE** | -83.5975 | 9.388383 | 1219 | Costa Rica |
| **QUEB** | -83.685 | 9.441727 | 1261 | Costa Rica |
| **GIGA** | -83.6347 | 9.455833 | 1407 | Costa Rica |
| **TOLO** | -83.7008 | 9.474174 | 1681 | Costa Rica |
| **SANM** | -83.7007 | 9.500052 | 1973 | Costa Rica |
| **BOQU** | -83.6923 | 9.484201 | 1993 | Costa Rica |
| **MILL** | -83.6867 | 9.561053 | 2710 | Costa Rica |
| **AMIG** | -83.7137 | 9.546838 | 2778 | Costa Rica |
| **NIMB** | -83.7413 | 9.565307 | 3107 | Costa Rica |
| **Mashpi_Capuchin** | -78.879 | 0.159003 | 774 | Ecuador |
| **MashpiLaguna** | -78.8702 | 0.170722 | 1020 | Ecuador |
| **UnPocoChoco** | -78.8409 | 0.051765 | 1046 | Ecuador |
| **Maquipucuna** | -78.6293 | 0.106333 | 1415 | Ecuador |
| **Sachatamia** | -78.7478 | -0.03683 | 1559 | Ecuador |
| **SantaLuciaLower** | -78.5995 | 0.11624 | 1776 | Ecuador |
| **LasGralarias** | -78.7347 | -0.009 | 1840 | Ecuador |
| **SantaLuciaUpper** | -78.5864 | 0.130211 | 2006 | Ecuador |
| **Alaspungo** | -78.6309 | 0.000904 | 2484 | Ecuador |
| **Verdecocha** | -78.5977 | -0.12051 | 3227 | Ecuador |
| **Yanacocha** | -78.5902 | -0.12118 | 3406 | Ecuador |

Table S2: Coefficients and associated standard errors of the (generalized) linear models used to describe elevation patterns in trait values and in the proportion of forbidden links

| Response variable | Explanatory variable | Estimate | Std. Error |
|---|---|---|---|
| **Trait values (link function: identity)** | Intercept (Ref: Brazil & birds) | 23.6976 | 3.292974 |
| | elevation (Ref: Brazil & birds)) | -0.00099 | 0.002676 |
| | Type (plants) | 13.88486 | 3.864534 |
| | Country (Costa Rica) | 3.428052 | 5.085071 |
| | Country (Ecuador) | -0.4746 | 4.639766 |
| | elevation:typeplants | -0.00213 | 0.003212 |
| | elevation:CountryCosta Rica | -0.00086 | 0.003554 |
| | elevation:CountryEcuador | 0.001643 | 0.003163 |
| | typeplants:CountryCosta Rica | 1.13005 | 5.917661 |
| | typeplants:CountryEcuador | -1.79614 | 5.320343 |
| | elevation:typeplants:CountryCosta Rica | -0.00488 | 0.004176 |
| | elevation:typeplants:CountryEcuador | -0.00201 | 0.003727 |
| **Proportion of forbidden links (link function: logit)** | Intercept (Ref: Brazil) | -1.00094 | 0.100588 |
| | elevation (Ref: Brazil) | -0.00017 | 7.65E-05 |
| | Country (Costa Rica) | -0.29056 | 0.184808 |
| | Country (Ecuador) | -0.69194 | 0.211812 |
| | elevation:CountryCosta Rica | -0.00035 | 0.000123 |
| | elevation:CountryEcuador | -5.55E-05 | 0.000124 |
